# Trade-offs between surviving and thriving: A careful balance of physiological limitations and reproductive effort under thermal stress

**DOI:** 10.1101/2025.11.07.687289

**Authors:** David L. Hubert, Ehren J. Bentz, Robert T. Mason

## Abstract

Balancing survival and reproduction presents a fundamental evolutionary challenge, especially in extreme and unpredictable environments. Thermoregulatory behavior in particular imposes a costly trade-off, as time spent maintaining optimal body temperature precludes other essential activities and forces individuals to balance competing selective pressures. By combining field observations, behavioral assays, and gene expression profiling of red-sided garter snakes (*Thamnophis sirtalis parietalis*), we describe how an ectothermic vertebrate navigates this trade-off in an extreme thermal environment through a combination of finely tuned mechanisms of behavioral and physiological plasticity. Snakes demonstrated remarkably consistent critical thermal limits, with minimal individual variation, suggesting a hard physiological constraint. Importantly, behavioral temperature thresholds for voluntary withdrawal and courtship cessation occurred within 5 °C of lethal, demonstrating that males routinely operate perilously close to lethal body temperatures in order to maximize reproductive opportunities. Context-dependent plasticity allows males to prioritize mating opportunities over thermoregulation until survival becomes an immediate threat. Molecular analyses of multiple tissues revealed rapid, tissue-specific transcriptomic responses activated within one hour of thermal challenge, well before critical limits were reached. Heat shock proteins were universally upregulated across all tissues under both heat and cold stress, suggesting anticipatory protective mechanisms that support risky behavioral decisions rather than simply responding to thermal damage. Tissue-specific patterns of gene expression reflected functional priorities with liver showing extensive metabolic flexibility, heart maintaining conservative stability, while brain and testis appeared to balance critical functions with stress responses. These findings demonstrate precise thermal reaction norms that integrate behavioral and molecular plasticity to maximize reproductive effort without compromising survival. The narrow margin between behavioral thresholds and lethal limits suggests this system may already be approaching evolutionary constraints. While phenotypic plasticity currently buffers populations against extreme thermal variability, it may paradoxically limit long-term adaptive potential as increasing environmental temperatures further narrow safety margins. Precise thermal decision-making, rapid physiological responses, and context-dependent behavioral switching suggest an evolved solution to the problem of balancing competing selective pressures.

## Introduction

Selective forces favoring either survival or reproduction are often placed in direct competition when finite time or energy must be allocated between the two. Thermoregulatory behavior in particular imposes a costly trade-off, as time spent maintaining optimal body temperature precludes other essential activities and forces individuals to balance competing selective pressures (Zera & Harshman, 2001). This dynamic varies across taxa resulting in diverse behavioral strategies (Huey & Slatkin, 1976; Liwanag, 2010; Van de Ven et al., 2020). For example, common wall lizards (*Podarcis muralis*) prioritize predator avoidance over thermoregulation under high predation risk (Amo et al., 2004), while climate-driven thermoregulatory shifts often come at the expense of growth and reproduction in other lizard species (Sinervo et al., 2010).

Optimizing survival and reproduction in extreme or variable thermal environments is particularly challenging and often results in costly trade-offs (Kingsolver & Huey, 1998). Many strategies exist for coping with these challenges, such as relying on precise behavioral timing to exploit narrow windows for foraging, growth, and reproduction in order to maximize fitness (Bogert, 1949; Besson & Cree, 2010). Common strategies include seasonal migration between mating and foraging grounds (King & Duvall, 1990; Madsen & Shine, 1996; Southwood & Avens, 2010), collective overwintering in hibernacula (Cowles, 1941; Ultsch, 2006; Huey et al., 2021), and large-scale reproductive aggregations (Garstka et al., 1982; Skinner et al., 2020). Although effective, each of these strategies imposes a trade-off between limited resources.

To mitigate the costs of trade-offs, many organisms demonstrate physiological or behavioral plasticity by dynamically regulating their phenotypes across environmental or behavioral contexts (Reznick & Endler, 1982; Pigliucci, 2001; Seebacher et al., 2014). Plasticity itself is an evolved trait measurable as reaction norms representing phenotypes that vary across contexts (Via & Lande, 1985; West-Eberhard, 2003; Dingemanse & Wolf, 2010). Behavioral plasticity can be observed in birds such as tree swallows (*Tachycineta bicolor*) and hooded warblers (*Setophaga citrina*) which adjust nesting, parental care, and foraging behavior in response to temperature variability (Ardia et al., 2006; Williams et al., 2025). Adaptive plasticity can buffer organisms against environmental conflict (Ghalambor et al., 2007; Seebacher et al., 2014). Yet plasticity is not without cost, as it can reduce physiological and metabolic performance (Seebacher & Franklin, 2005; Glanville & Seebacher, 2006), alter energy allocation (Kearney et al., 2009), affect gestation and reproductive outcomes (Lourdais et al., 2002), and reshape temporal resource use (Beever et al., 2017).

These challenges are further amplified in a rapidly changing climate, as global temperatures rise and thermal environments become increasingly unpredictable (Parmesan, 2006; Pimm et al., 1995; Walther et al., 2002; Parmesan & Yohe, 2003; Murphy-Klassen et al., 2005; Ceballos et al., 2017; Bellard et al., 2012; IPCC, 2021). Global warming results in increasingly unpredictable environmental temperatures and narrows thermal safety margins (Parmesan & Yohe, 2003; Walther et al., 2002; Parmesan, 2006; IPCC, 2021). Population-level impacts are evident, such as extinctions in lizards (Sinervo et al., 2010) and widespread vertebrate declines (Ceballos et al., 2017), suggesting that failure to maintain the balance between survival and reproduction is becoming increasingly costly and difficult to mitigate. This is especially true for ectothermic vertebrates which rely entirely on the external environment to regulate body temperature and metabolic rate (Besson & Cree, 2010; Huey et al., 2021).

Red-sided garter snakes (*Thamnophis sirtalis parietalis*) provide an exceptional system in which to examine the intersection of thermal challenge, phenotypic plasticity, and reproductive trade-offs. This common, broadly distributed species ranges from Texas to the Northwest Territories of Canada (Aleksiuk & Stewart, 1971). The most intensively studied population inhabits the Interlake region of Manitoba, where snakes display striking seasonal migration patterns and extreme cold tolerance near the northern edge of their range (Gregory, 1974; Garstka et al., 1982; Shine et al., 2000; Lutterschmidt et al., 2006).

Winter temperatures in this region regularly fall below −40 °C, with persistent snow cover and freezing conditions lasting up to eight months (Environment Canada, 2014; Bush & Lemmen, 2019). The onset of spring initiates a dramatic behavioral and thermal transition in which individuals experience an extremely rapid rise in body temperature as they emerge from hibernacula at near-freezing subterranean temperatures (<1 °C) into surface conditions that may exceed 40 °C (Aleksiuk & Gregory, 1974; Whittier et al., 1987). Thermal conditions remain unpredictable throughout the spring mating season, with ambient temperatures often fluctuating from below freezing to greater than 40 °C within a single day (Environment Canada, 2014; Bush & Lemmen, 2019). Unsurprisingly, these rapidly changing thermal conditions can easily become lethal unless snakes continuously utilize behavioral thermoregulation.

Despite these risks, male snakes remain near the den for 4-6 weeks of intense reproductive activity. They do not feed during this time, relying entirely on energy reserves stored the previous summer and fall (O’Donnell et al., 2004). Mating aggregations often involve more than 10,000 snakes clustered within meters of the den (Garstka et al., 1982; Shine et al., 2001), where males must withstand thermal extremes, avoid predation, and compete intensely for access to females. Although costly and dangerous, males are obliged to remain within these conditions as long as possible to maximize reproductive opportunity.

The convergence of energetic stress, physiological constraints, and competition during this brief but intense mating period highlights selective pressures favoring traits that maximize both survival and reproductive success. Male red-sided garter snakes must balance thermoregulation, predator avoidance, and prolonged reproductive effort, often while operating near their thermal limits.

Because coping with an extreme and unpredictable thermal environment constitutes one of the greatest challenges for male snakes at the den, we closely examined their thermoregulatory behavior and thermal physiology to better understand how ectothermic vertebrates cope with a trade-off between survival and reproduction in harsh and variable environments.

This study included four specific aims: 1) Characterize the thermoregulatory behavior and typical body temperatures of freely behaving snakes in their natural habitat. 2) Quantify the absolute physiological thermal limits of the species under controlled laboratory conditions where thermoregulation is not possible. 3) Identify temperature thresholds and behavioral contexts under which snakes adjust their behavior to avoid lethal exposure. 4) Examine transcriptional responses to acute thermal stress near the upper and lower thermal tolerance limits using high-throughput RNA sequencing.

To address these objectives, we used an integrative approach combining field observations, laboratory experiments, and molecular profiling to provide a comprehensive understanding of the strategies male red-sided garter snakes use to optimize survival and reproductive success in extreme thermal environments.

## Materials and Methods

### Animal collection, husbandry, and ethics review

All procedures were conducted using Red-sided garter snakes collected from the natural den site near the town of Inwood, Manitoba, Canada during the peak of the spring mating period (April-May). Animals were maintained in outdoor nylon cloth arenas (1×1×1 m) for the duration of experiments up to a maximum of 14 days and housed in groups with males and females in separate enclosures. Enrichment items including hide boxes and stones to facilitate shedding were available at all times. Water was provided *ad libitum*, and no food was offered, because *T. s. parietalis* is aphageous during the spring mating period (O’Donnell et al., 2004). All animals included in the study were either returned to the site of capture or euthanized for tissue collection in accordance with approved ethics protocols. All procedures were approved by the Oregon State University Institutional Animal Care and Use Committee under protocol 4818. Field research in Canada was conducted under the authority of Manitoba Wildlife Scientific Permit No. WB16264.

### Body temperatures of freely-behaving wild snakes at the den

To determine the internal body temperature of wild male garter snakes, temperatures were measured at midday on a cool day (T_cool_) with an air temperature of 11.50 °C (n=45), a warm day (T_warm_) with an air temperature of 17.50 °C (n=42), and a hot day (T_hot_) with an air temperature of 22.00 °C (n=20). All body temperature data collected were obtained via cloacal probe using a calibrated Fluke 51-Series II thermometer with a k-type thermocouple (Fluke Corp., Everett, WA). Animals selected for temperature recordings were selected based on the most commonly observed behavior at that time. Cool- and warm-day individuals were chosen if they exhibited typical courtship behavior. Because no courtship behavior was observed at the time of data collection during the hot day, body temperatures were recorded from inactive animals (hereby referred to as ‘loafing’).

### Behavioral thermal maxima

Two experimental assays were designed to establish behavioral maximum temperature thresholds. The voluntary maximum temperature (BT_vol_; the body temperature threshold at which a snake seeks to escape thermal conditions in the absence of potential mating opportunities), and the courtship maximum temperature (BT_court_; the body temperature threshold at which a snake would cease courtship behavior to escape high temperatures).

The voluntary maximum temperature was measured by placing individual snakes (n=20) into a high temperature enclosure (Figure 1A) without the ability to behaviorally thermoregulate until specific behavioral cues were observed that indicated an active attempt to escape the thermal conditions. The high temperature enclosure consisted of an inverted 1.2 L rectangular clear Pyrex bowl on an electric heating pad (Jumpstart, 17 W, 120 V) in a 10-gallon glass aquarium, with a ceramic heat emitter (250 W, 110 V) fixed 17 cm above the top of the inverted bowl. The heating pad and the ceramic heat emitter were each controlled by a separate digital temperature controller (Bayite BTC201) and maintained an air temperature of 43.00 °C (± 0.20 °C). Briefly, snakes would settle into a stationary position after approximately 2 minutes of mild exploration activity and remain inactive for approximately 8 minutes, at which point they became very active, trying to escape the enclosure. Once a snake exhibited this escape behavior it was removed from the enclosure and their body temperature was immediately measured and recorded and the animal was returned to normal thermal conditions.

**Figure 1.**
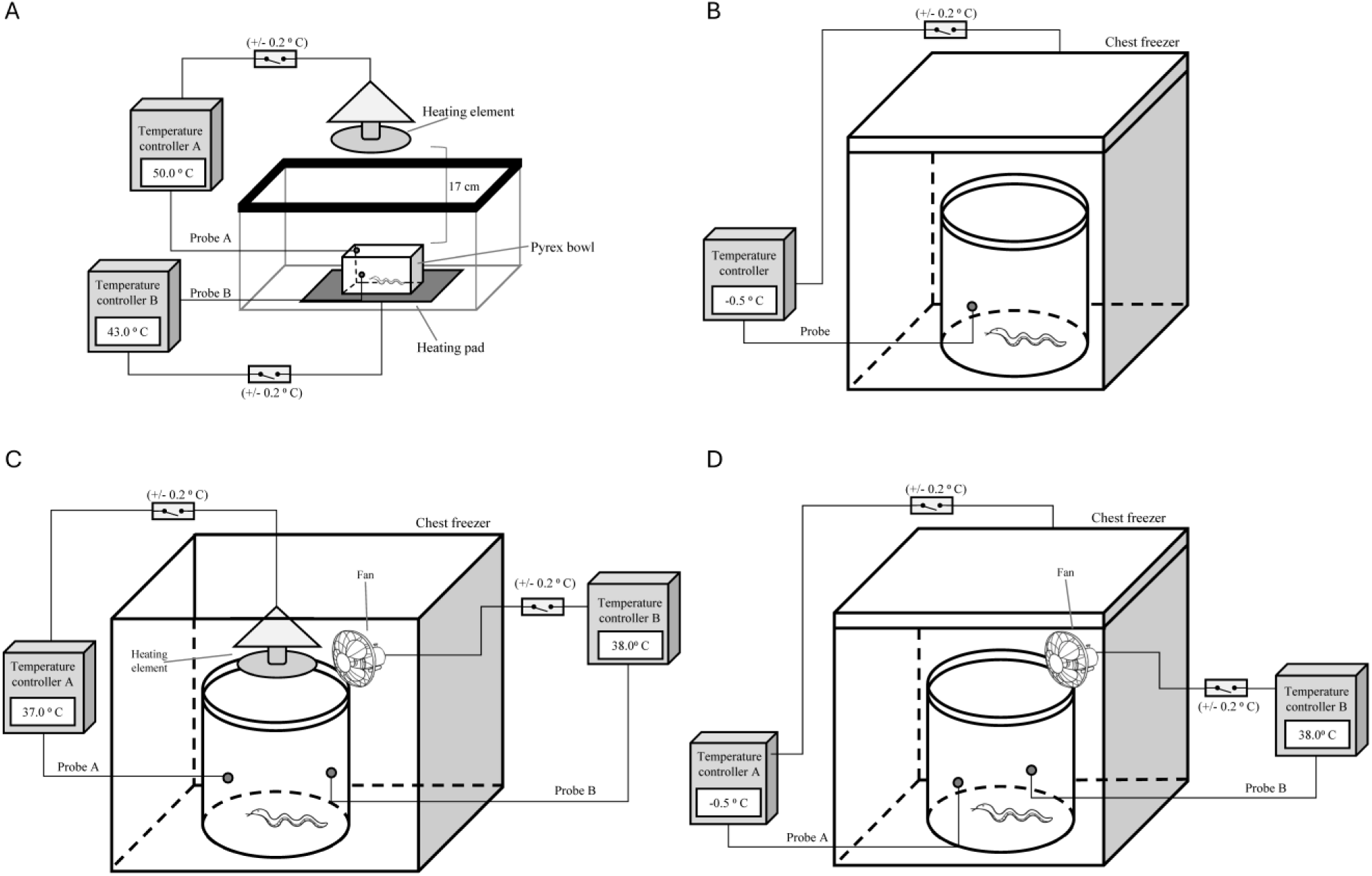
Thermal chamber diagrams. A) High temperature enclosure maintained a mean air temperature of 43.00 ± 0.20°C and was used for both the behavioral maximum (BT_max_) and the critical thermal maximum (CT_max_) experiments. B) Low temperature enclosure maintained a mean air temperature of -0.50 ± 0.20°C and was used for the CT_min_ experiment. C) Acute heat stress chamber maintained a mean air temperature of 39.00 ± 0.50°C and was used for the transcriptional response to acute heat stress experiment. D) Acute cold stress chamber maintained a mean air temperature of 0.00 ± 1.00 °C and was used for the transcriptional response to acute cold stress experiment.

**Figure 2.**
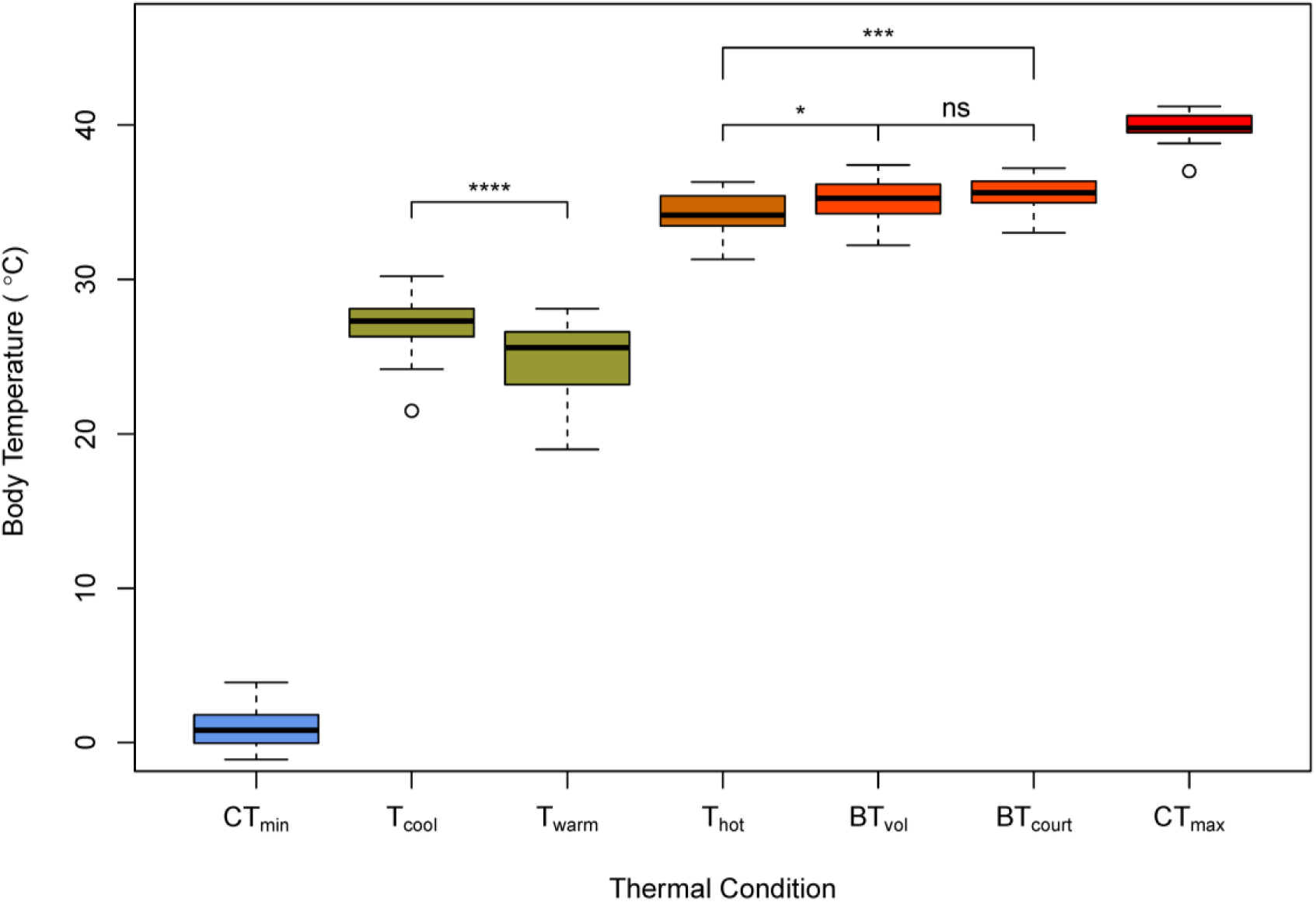
Internal body temperatures of male Red-sided garter snakes. Temperatures from free ranging snakes on a cool day (T_cool_), a warm day (T_warm_), and a hot day (T_hot_). Experimentally derived temperatures from snakes at physiological maximums, critical thermal maximum (CT_max_) and minimum (CT_min_), and behavioral maximums, voluntary maximum (BT_vol_), and courtship maximum (CT_court_). All temperatures were obtained vial cloacal probe. Physiological maxima was determined by loss of righting reflex, and behavioral maxima was determined through behavioral change (methods). Significance for select comparisons determined through Welch two sample t-test.

The maximum courtship temperature was measured by conducting courtship behavioral assays while the environmental temperature was increased until courtship behavior ceased. Red-sided garter snakes (n=25 males & 10 females) were randomly sorted into 5 groups consisting of 5 dorsally marked male snakes and 2 female snakes with their cloaca obstructed to prevent copulation during the experiment as described in Friesen et al. (2017). Snakes were introduced into a warmed (∼13.00 °C) arena (an opaque, 0.5 m diameter, 1 m tall cylindrical nylon enclosure) with a heat lamp positioned above and a heating pad beneath it. No water, substrate, or hides were provided during the courtship behavioral assay to minimize any additional behavioral thermoregulation. The snakes were allowed 5 minutes to acclimate to the warmed arena and begin courtship behavior, after which, the heat lamp was turned on to raise the ambient temperature of the arena. Behavior was continually monitored and scored using the ethogram of male courtship behavior (Table 1). Any male that failed to attain a courtship score of 4 within the 5-minute acclimation time was removed from the trial. Any of the remaining males that stopped courtship behavior (courtship score ≤1) were manually prodded, and if they resumed courtship behavior they were allowed to remain in the trial, if they did not resume courtship behavior they were removed, and their body temperature was immediately recorded. The trial was continued until all males ceased courtship behavior.

**Table 1:** Ethogram of male courtship behavior.

| **Courtship score** | **Description of male behavior** |
| --- | --- |
| 0 | No courtship behavior exhibited |
| 1 | Investigates female |
| 2 | Chin-rubs female and/or rapidly tongue-flicks female's skin |
| 3 | Aligns body with female |
| 4 | Actively tail-searches female, attempts cloacal apposition; possible caudocephalic waves |
| 5 | Copulation or attempted intromission |

(Modified from Friesen et al., 2017; Moore et al., 2000; and Crews 1984)

### Critical thermal minima and maxima

The critical thermal maximum (CT_max_) and the critical thermal minimum (CT_min_) represent specific temperatures defining the absolute boundaries of an organism’s thermal tolerance to describe the complete range of survivable thermal conditions (Lutterschmidt and Hutchison 1997; Spellerberg 1972; Ernst et al. 2014). In this study, we define CT_max_ and CT_min_ as the minimum or maximum body temperatures at which a snake was observed to lose locomotory coordination resulting in the loss of righting response (the ability to right itself when placed on its back). This is an effective indicator of physiological thermal limits as it is the point at which they can no longer escape from thermal conditions that would otherwise lead to death (Spellerberg 1972).

Both CT_min_ and CT_max_ were experimentally determined under tightly controlled conditions. Male snakes (n=25 per treatment) were placed individually into either a high temperature enclosure (Figure 1A) or a low temperature enclosure (Figure 1B). The low temperature enclosure consisted of a five-gallon bucket placed inside of a small insulated refrigerated enclosure set to -0.50 (± 0.20) °C. The temperature of the enclosure was controlled by a Bayite BTC201 digital temperature controller with a probe set to measure air temperature where the animal was located. At the time each snake was introduced to the low temperature enclosure, surface temperature of the bottom of the inside of the bucket was recorded with a Ryobi Tek4 IR thermometer and had a mean temperature of -4.19 (± 0.50 °C).

The behavior of each animal was continually monitored while body temperatures were allowed to rapidly change until a behavioral cue was exhibited, then the snake was tested for loss of locomotory coordination, as measured by loss of righting response (Spellerberg, 1972).

Briefly, the animal was placed on its dorsal side and allowed to return to its ventral side without assistance. Once an animal could not right itself, its body temperature was immediately measured and recorded, and the snake was returned to normal thermal conditions.

The behavioral cues differed between treatments. For the high-temperature treatment, snakes would settle into a stationary position after approximately 2 minutes of mild exploration activity and remain inactive for approximately 8 minutes, at which point they became very active, trying to escape the enclosure (BT_vol_). The escape behavior was followed by a second inactive period, often marked a yawning behavior (gaping of the mouth) consistent with previously reported behavioral observations made near critical thermal temperatures in snakes (Cowles and Bogert 1944). This second inactive period was determined to be an accurate behavioral signal that the animal had lost locomotory coordination (CT_max_), as measured by loss of righting reflex.

For the low temperature treatment, snakes would settle into a stationary coiled position after approximately 6-8 minutes of mild exploration activity. This inactivity and coiled posture was determined to be an accurate behavioral signal that the animal had lost locomotory coordination (CT_min_), as measured by loss of righting reflex.

### Transcriptome sequencing, assembly, and annotation

Immediately prior to tissue collection snakes were given a subcutaneous injection of a lethal overdose methohexital sodium (Brevital™) (0.005 mL/g of body mass, 1% solution), then returned to conditions for approximately 2-3 minutes to allow the anesthetic to take effect. Euthanasia was confirmed by performing a snout-tap reflex response assay, testing for neural function by gently tapping the tip of the snout to elicit a reflexive spasm. Once the animal no longer exhibited the reflexive spasm, tissues were surgically removed and immediately placed in >10 volumes of RNAlater and stored at 4 °C for 24 hours then transferred to -20 °C until RNA extraction.

A sample of approximately 10 mg of tissue was mechanically homogenized for each replicate and RNA extraction from homogenized tissue was conducted using the Omega Bio-Tek E.Z.N.A® HP total RNA kit and following the standard protocol. RNA quality was evaluated and visualized using gel electrophoresis and concentration estimates were made for each sample using absorbance assays.

Total RNA was extracted from a total of 16 samples comprised of 6 tissues: whole brains (n=1 male & 1 female), liver (n=1 male & 1 female), kidney (n=1 male & 1 female), testes (n=2 male), Harderian gland pairs (n=2 male & 2 female), and vomeronasal organs (n=2 male & 2 female). An equal quantity of total RNA (500ng) from each sample was added to a sample pool. The pool was diluted to a concentration of 111ng/μl, then poly-A enriched and reverse transcribed to cDNA using Bioline Tetro reverse transcriptase and custom primers (Table S1). An abundance-normalized cDNA library was prepared to avoid over-sequencing of abundant transcripts and to improve representation of rare transcripts (Meyer et al., 2009; Kitchen et al., 2015). The completed library was submitted to the Genomics & Cell Characterization Core Facility (University of Oregon, Eugene, USA) and 150 bp paired-end reads were generated using 2 lanes on the Illumina MiSeq platform.

Approximately 82M read-pairs were quality filtered and removed from the dataset if either read from a pair contained ≥ 20bp with a Quality Value ≤ 20, ≥50 homologous repeats, or ≥15bp aligning to Illumina adapter sequences. Reads passing quality filters were examined using FastQC v0.11.3 (Andrews, 2010). Paired-end reads with valid mates assembled using Trinity v2.1.1 (Grabherr et al., 2011; Haas et al., 2013). Contigs of 200bp or larger were retained in the final assembly. To remove isoform redundancy, the assembly was filtered to retain the trinity sub-component with the longest open reading frame resulting in a final assembly containing 83,394 transcripts.

Transcripts were annotated using the Blast2GO sequence annotation suite v6.0.3 (Gotz et al., 2008). Transcripts were searched using BLASTx against all vertebrate sequences obtained from the NCBI non-redundant protein database. The top 20 hits for each sequence were retrieved and filtered to remove low quality or ambiguous matches (e-value ≤ 0.001, HSP length ≤ 30aa, and unidentifiable or non-specific matches including terms “Whole Genome Shotgun”, “Unidentified”, “Unknown”, “Hypothetical Protein”, or “RIKEN”). Matches passing all filters were used to assign NCBI accession numbers and protein names to transcripts using the Blast2GO description annotator with evidence codes equally weighted. Protein IDs were mapped to the Gene Ontology (GO) database (Gene Ontology Consortium, 2023; Ashburner et al., 2000) to assign functional annotation terms. Best matches were assigned, and final GO annotations were organized and redundancy removed according to the go.obo 01-2025 GO hierarchy definitions file.

### Transcriptional response to acute thermal stress

To evaluate the transcriptional response to acute thermal stress, male garter snakes were exposed to either acute heat or cold thermal stress conditions designed to maintain specific internal body temperatures for the duration of 1 hour. Thermal stress conditions were controlled such that sustained internal body temperatures would remain near, but not exceeding, the critical thermal limits previously identified during Aim 2 of this study. A control group was similarly housed and handled but were not exposed to thermal stress conditions, and instead maintained body temperatures of approximately 20 °C for the duration of the experiment. Snakes were placed into either the acute heat stress chamber (n = 3) (Figure 1C), the acute cold stress chamber (n = 3) (Figure 1D), or maintained at normal environmental conditions (n = 3). The acute heat stress chamber consisted of a food grade plastic five-gallon bucket placed inside of a small, insulated enclosure to provide a thermally stable environment. The temperature inside the apparatus was controlled by a pair of digital temperature controllers (Bayite BTC201) with both probes set to measure the air temperature inside the bucket. One controlled a ceramic heat emitter (250W, 110V) positioned above the bucket and heating pads placed below and set to 39.0 (± 0.5) °C. The other digital temperature controller controlled a fan that was positioned above the bucket and was set to circulate air when the temperature rose above 39.5 °C, resulting in a slight drop in temperature. This provided a consistent average air temperature of 39.0 (± 0.5) °C and resulted in an average sustained body temperature of 35.8 °C (± 0.9 °C) for the duration. The acute cold stress chamber consisted of a five-gallon bucket that was placed inside of an insulated refrigerated enclosure set to 0.0 (± 0.5) °C. Temperature was controlled by a pair of digital temperature controllers (Bayite BTC201) with both probes set to measure the air temperature inside the bucket, one controlling the freezer and one controlling a fan. The fan was positioned above the bucket and was set to blow air across the top when the temperature fell below -0.5 °C, which caused a slight rise in temperature. This resulted in an average air temperature of 0.0 (± 1.0) °C and an average sustained body temperature of 2.0 °C (± 1.0 °C) for the duration.

After 1 hour in treatment conditions, final body temperatures were recorded. Animals were euthanized for tissue collection, including brain, heart, liver, and testis. Tissue samples were collected within 5 minutes of leaving treatment conditions to preserve target transcriptional response.

Three biological replicates were sequenced from each treatment-tissue condition for a total of 36 samples. Total RNA (1.3 µg per sample) was poly-A selected and used to produce individual sequencing libraries using the Tag-seq protocol (Meyer et al., 2011). Completed libraries were pooled in equal quantities and submitted to Oregon Health & Science University, Massively Parallel Sequencing Shared Resource (MPSSR) and 100bp single-end reads were generated using one lane on the Illumina HiSeq 2500 platform. Resulting raw read quality was analyzed using FastQC (version 0.11.5) (Andrews 2010).

Raw reads were trimmed of leading adapter sequence, then quality filtered to remove reads with ≥ 20bp with a Quality Value ≤ 20, ≥ 10bp homologous repeats, or ≥ 12bp aligning to Illumina adapter sequences. Reads passing all filters were mapped to the multi-tissue annotated reference transcriptome using the gmapper tool from the SHort Read Mapping Program (SHRiMP2) with the following options: --qv-offset 33 -Q --strata -o 3 - N 4 -K 10000 -L (David et al., 2011; Rumble et al., 2009).

Differential expression analysis was conducted in R using the DESeq2 package (Love et al., 2014). Transcripts were pre-filtered to remove very low abundance transcripts with a mean of fewer than 3 mappings within each condition. Within each of the four tissues, pairwise Wald tests were used to independently contrast conditions: 1) heat stress vs. control, and 2) cold stress vs control. Raw p-values were adjusted for multiple testing using the Benjamini-Hochberg false discovery rate (FDR) (Benjamini and Hochberg, 1995). Transcripts with an adj. p-value < 0.05 were considered significantly differentially expressed between conditions.

Functional enrichment analyses were performed to identify Gene Ontology (GO) terms enriched in genes either up- or down-regulated in response to heat stress or cold stress. Enrichment analyses were performed. Transcripts were ordered using signed p-values calculated from DESeq2 results, and functional enrichment was performed using bi-directional Mann-Whitney U tests to identify GO terms significantly enriched in up- or down-regulated transcripts. Raw p-values for each GO term were FDR corrected and GO terms with an adjusted p-value of < 0.05 were considered significantly enriched.

## Results

### Body temperatures of freely-behaving wild snakes at the den site

Active snakes showed significantly different mean body temperatures between cool and warm days, averaging 27.21 ± 0.23 °C (mean ± SE) on cool days and 24.98 ± 0.34 °C (mean ± SE) on warm days (Welch’s t-test, p = 8.0 × 10⁻⁷). Inactive snakes on hot days had a higher mean body temperature of 34.09 ± 0.34 °C (mean ± SE) (Table 2, Table S2).

**Table 2:** Internal body temperatures collected in the field and experimentally.

| **Thermal condition** | **Mean (95% CI) (°C)** | **Range (°C)** | **n** |
| --- | --- | --- | --- |
| T<sub>warm</sub> | 24.98 (24.30 - 25.70) | 19.00 – 28.10 | 42 |
| T<sub>cool</sub> | 27.21 (26.80 - 27.70) | 21.50 – 30.20 | 45 |
| T<sub>hot</sub> | 34.09 (33.40 - 34.80) | 31.30 – 36.30 | 20 |
| BT<sub>court</sub> | 35.55 (34.90 - 36.20) | 33.00 – 37.20 | 16 |
| BT<sub>vol</sub> | 35.12 (34.40 - 35.80) | 32.2 – 37.4 | 20 |
| CT<sub>max</sub> | 39.90 (39.50 - 40.30) | 37.00 - 41.20 | 25 |
| CT<sub>min</sub> | 0.97 (0.46 - 1.49) | -1.10 - 3.90 | 25 |

### Behavioral thermal maxima

Behavioral maximum temperatures did not differ significantly between contexts, with a mean voluntary maximum of 35.12 ± 0.34 °C (mean ± SE) and a mean courtship maximum of 35.55 ± 0.28 °C (mean ± SE) (Welch’s t-test, p = 0.34) (Table 2, Table S2). Both were significantly higher than the mean body temperatures of inactive snakes on hot days (Welch’s t-test, p = 0.04 and p = 2.4 × 10⁻³).

### Critical thermal maximum and minimum

Loss-of-righting response assays revealed thermal tolerance limits with narrow confidence intervals (< 1 °C). Mean CT_max_ was 39.90 ± 0.19 °C (mean ± SE), ranging from 37.00–41.20 °C (mean ± SE) across individuals, while mean CT_min_ was 0.97 ± 0.25 °C (mean ± SE), ranging from –1.10–3.90 °C (Table 2, Table S2). Heating rates averaged 2.60 ± 0.20 °C (mean ± SE) per minute (n = 45) across both CT_max_ and behavioral assays, substantially higher than the 0.83 °C per minute reported by Shine et al. (2000).

### Transcriptome assembly and annotation

After filtering, 43.4 M read pairs produced an assembly of 83,394 transcripts (min = 201 bp, max = 14,870 bp, mean = 716.69 bp, median = 441 bp, N50 = 1,015 bp across 15,600 sequences; GC content = 40.11%). BLASTx annotation identified 33,148 transcripts with matches, of which 21,070 were assigned at least one GO term. Hidden Markov models (Pfam) yielded 16,866 annotations, and 792,191 potential ORFs ≥ 20 aa were predicted.

### Transcriptional response to acute thermal stress

Illumina HiSeq 2500 generated ∼145.4 M 100 bp reads (∼4 M per sample). After filtering, 97.3 M reads remained (∼2.7 M per sample). One outlier was removed (Grubbs test, p = 8.8 × 10⁻⁶), leaving 96.9 M reads (∼2.8 M per sample). Mapping yielded 94.9 M aligned reads, covering 55,220 transcripts, with ∼15,364 transcripts per sample after filtering (range 11,572– 19,583).

### Differential Gene Expression

Differential expression analysis identified several transcripts responsive to acute thermal stress across tissues. After filtering out low-abundance transcripts (< 3 counts per condition), Wald test comparisons were performed in DESeq2 with false discovery rate (FDR) correction. Using a significance threshold of FDR < 0.05, we detected 337 (265 unique) transcripts that were significantly differentially expressed across tissues and treatments (Table S3 and Figure 3). Liver under hot conditions showed the highest response compared to the control condition (100 DE transcripts: 61 upregulated, 39 downregulated), while overall, heart tissue showed the fewest (47 total). Heat stress conditions induced the greatest transcriptional response, with 204 total significant DE transcripts (111 upregulated and 93 downregulated) relative to the control condition, while the cold stress response yielded a total 133 significant DE transcripts (70 upregulated and 63 downregulated) relative to the control condition. We see several of the most significant differentially expressed genes involve a response to thermal stress and maintaining protein homeostasis in both the heat and cold shock conditions (Figure 4).

**Figure 3.**
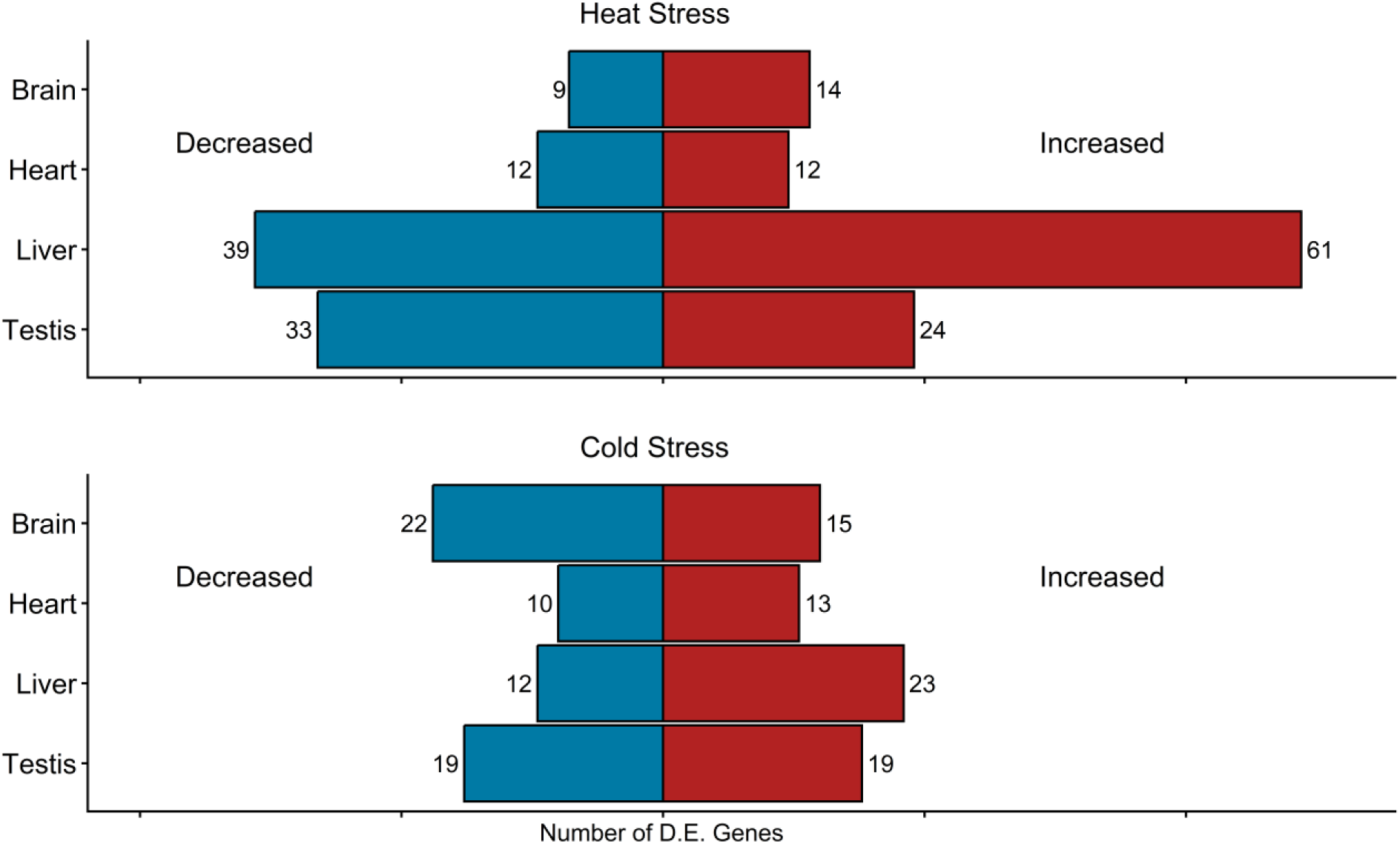
Numbers of significantly differentially expressed (D.E.) genes. identified by DESeq (FDR < 0.05) in four tissues (brain, heart, liver, and testis) under heat stress (top) and cold stress (bottom). Bars indicate genes with decreased (left) or increased (right) expression relative to control conditions. Heat stress elicited the strongest transcriptional response in the liver, while brain and testis showed more moderate changes. Cold stress responses were more balanced across tissues, with both increases and decreases observed.

**Figure 4.**
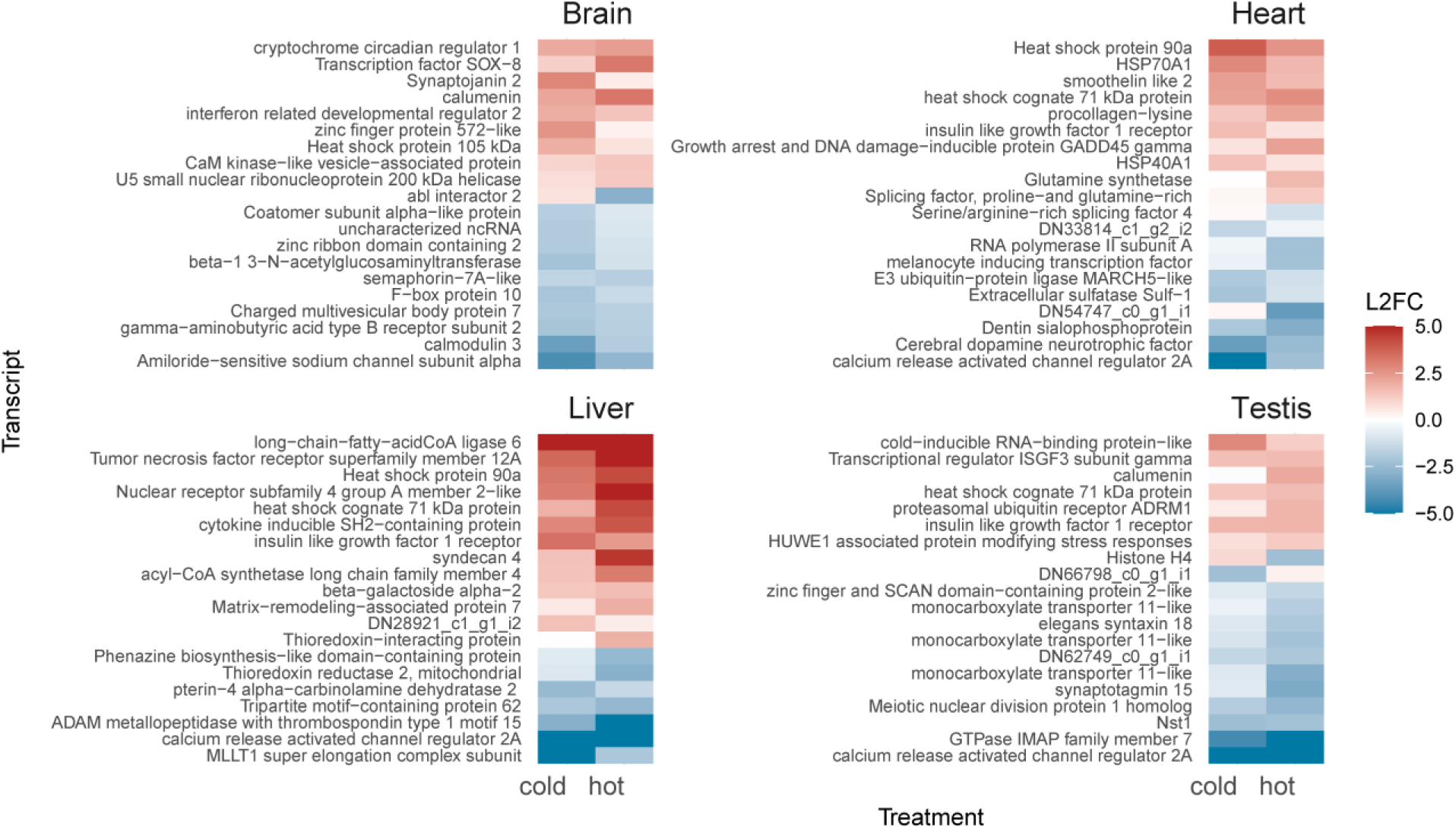
Heatmap of significantly differentially expressed genes. (FDR < 0.05) in four tissues (brain, heart, liver, and testis) under cold and heat stress relative to control conditions. Expression values are shown as log2 fold change (L2FC), with red indicating increased expression and blue indicating decreased expression. Each panel highlights the strongest transcriptional responses within that tissue, including induction of canonical stress response genes (e.g., heat shock proteins) as well as tissue-specific regulatory factors.

### Functional Enrichment

Functional enrichment analyses revealed distinct tissue- and temperature-specific transcriptional responses to thermal stress (Figure 5). We identified a total of 101 enriched GO terms: 30 in the brain, 11 in the heart, 43 in the liver, and 17 in testis tissue (Mann-Whitney U test, adj. p-value < 0.05; Table S4).

**Figure 5.**
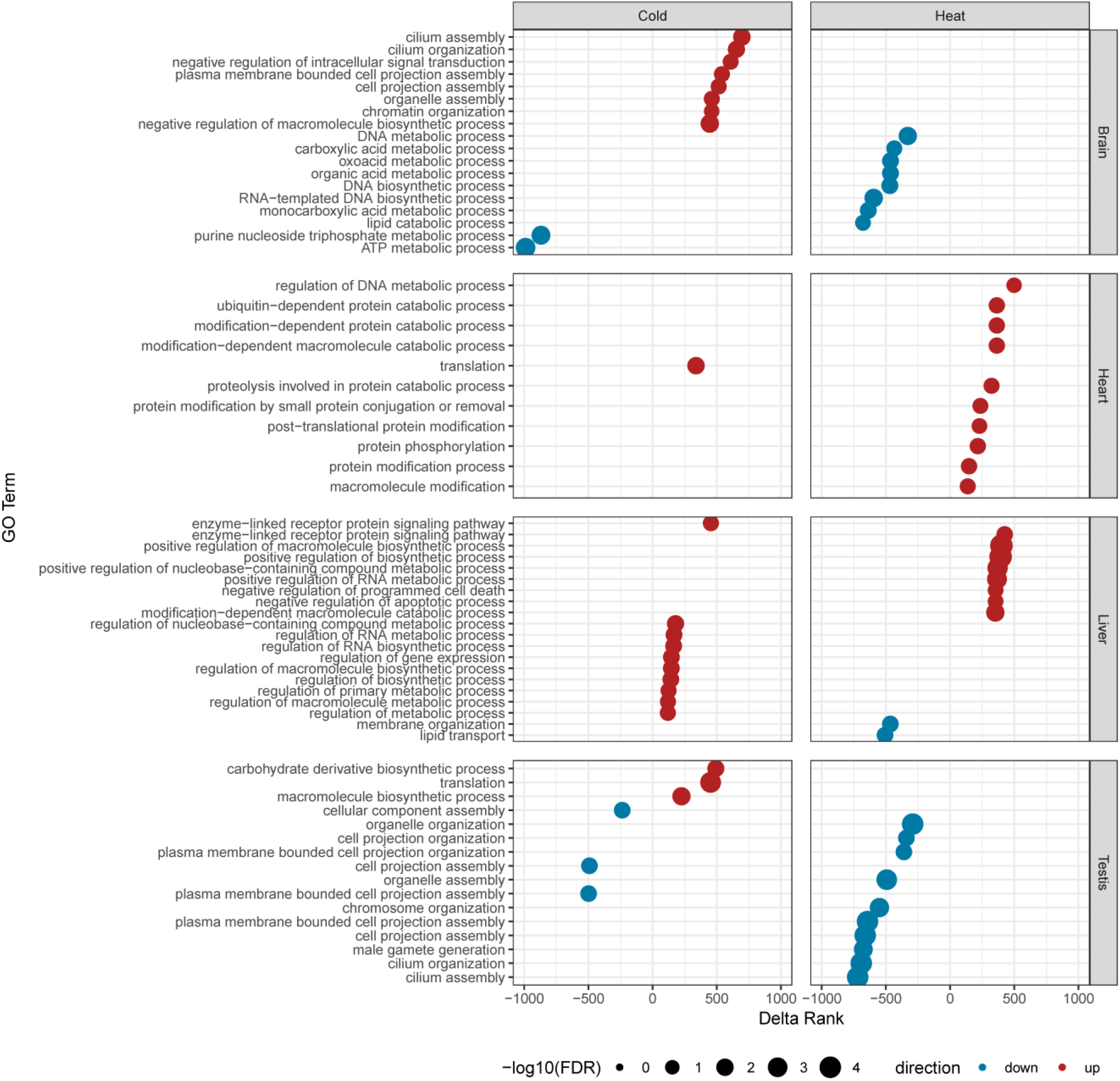
Gene Ontology enrichment analysis results by treatment and transcriptional regulation. Top 10 GO terms for each tissue and treatment (10 smallest FDR values that are <0.05). Color represents direction of regulation relative to control condition, and the delta rank represents the magnitude of that difference in expression level. Dot size represents significance level, with larger dots indicating more significant FDR values.

In the brain, genes downregulated in response to heat stress were enriched for multiple metabolic processes, and RNA and DNA biosynthesis. Genes downregulated in the brain in response to cold stress were enriched for several metabolic processes, whereas genes upregulated in response to cold stress were enriched for structural functions, chromatin organization, and negative regulation of metabolic processes.

In the heart, genes upregulated in response to heat stress were enriched for protein catabolic processes, and genes upregulated in response to cold stress were enriched for translation.

In the liver, genes upregulated in response to heat stress were enriched for many functions including metabolism, catabolism processes, cell signaling and RNA and protein processing, whereas genes downregulated in response to heat stress were enriched for membrane organization and lipid transport. Genes upregulated in the liver in response to cold stress were enriched for gene expression functions, transcription, and RNA processing.

In testis tissue, genes downregulated in response to heat stress were enriched for cellular structure and sperm production. Genes downregulated in the testis in response to cold stress were enriched for cellular structure, and upregulated genes were enriched for carbohydrate metabolism and translation.

## Discussion

### Physiological thermal limits and behavioral thresholds

We observed remarkably little variation between individual critical thermal limits at both the upper and lower extremes of their physiological range. This narrow range is consistent with molecular mechanisms involving thermal instability of critical proteins or enzymatic systems that fail at extreme temperatures (Licht 1964; Gangloff & Telemeco, 2018; Bennett et al., 2021). These effects were observed to be only temporary, as righting responses were immediately restored as body temperatures returned to normal, indicating that the failure was due to transient protein instability rather than permanent denaturation (Somero 1995; Williams et al., 2016).

The thresholds at which individuals modified their behavior in response to rising body temperatures were only slightly less variable than the hard physiological limits (BT_vol_: 35.12°C ± 0.34°C (mean ± SE), range 33.20-37.00; BT_court_: 35.55 ± 0.28°C (mean ± SE), range 33.80-37.20°C). This narrow range suggests that individual snakes actively monitor minute changes in body temperature enabling them to make decisions regarding thermal risk and, more importantly, that those decisions varied across different contexts.

The proximity of behavioral thermal maxima to the critical thermal maximum was only 4.35°C. Field observations show that inactive snakes on hot days maintained body temperatures of 34.10°C, which is significantly less than both behavioral maxima yet still approaching dangerous thresholds, demonstrating that snakes routinely operate surprisingly close to lethal body temperatures.

The observed heating rate of 2.60°C/min is more than 3x previously reported rates (Shine et al., 2000). Although the ability to tolerate rapid changes to body temperature may be advantageous for quickly achieving optimal temperatures for courtship, rapid heating can also cause snakes to quickly approach lethal temperatures and therefore require careful behavioral thermoregulation to stay within critical limits.

### Behavioral plasticity maximizes reproductive opportunities

Male snakes demonstrate context-dependent behavioral plasticity that prioritizes reproduction over thermal safety until rising body temperatures cross a threshold within just a few degrees of becoming lethal. The courtship maximum was greater than body temperatures of inactive snakes on hot days, suggesting that males are more willing to approach the critical maximum if mating opportunities are available. Furthermore, the lack of a difference between the voluntary maximum and courtship maximum indicates that males will maintain courtship behavior until thermal stress triggers an escape response, demonstrating that males quite literally continue to exploit potential mating opportunities until they must flee to survive.

Red-sided garter snakes represent an archetypal seasonal explosive breeding species in which large mating aggregations create intense competition for mating opportunities (Erisman et al., 2017; Arak, 1983; Bradbury & Gibson, 1983). Because the brief 4-6 week mating period represents the sole annual reproductive opportunity for the entire population (Garstka et al., 1982), males cannot wait for conditions to improve nor can they feed to replenish energy reserves (O’Donnell et al., 2004). Every aspect of this reproductive system drives males toward accepting greater thermal risk to be present and active in order to pursue potential mating opportunities. In this context, making thermally risky decisions not only make evolutionary sense, but is likely obligatory, as an individual that abandons courtship prematurely due to thermal stress will be thoroughly outcompeted by other males.

The extremely consistent critical thermal maximum in this species likely facilitates this highly refined thermally mediated behavioral plasticity, as the existence of a very consistent lethal maximum allows an individual to be acutely aware of his thermal risk and allow only a minimal temperature buffer to escape potentially lethal thermal conditions.

### Transcriptional responses represent physiological plasticity supporting thermally risky behavior

We observed transcriptional responses within one hour of exposure to both high and low temperatures. These responses were activated at body temperatures below behavioral switching thresholds and well before critical thermal limits were reached, indicating that molecular protective mechanisms operate in anticipation of, rather than in response to, thermal damage.

Specifically, transcriptional responses were already active within the range of body temperatures where snakes were observed actively engaging in courtship behavior. Though we can conclude that thermal stress results in a plastic response observable as tissue-specific or universal changes to gene expression, we may only speculate about the exact physiological functions or evolutionary implications of these findings.

Expression of genes encoding heat shock proteins was increased in response to both high and low body temperatures in all tissues, suggesting that thermal protection was universally increased in response to thermal challenge, regardless of direction or tissue. This pattern of expression may be due to the thermal environment at the den, in which temperatures may fluctuate from below freezing to lethally high within the same day, requiring molecular thermal protective mechanisms to be universal rather than specific to either high or low temperatures.

However, this pattern may also be a more broadly conserved evolutionary mechanism, as HSPs are commonly known to act as chaperones to confer protection to both heat and cold damage (Howarth & Ougham 1993; Goto et al., 1998; Zhao et al., 2014; Lindquist & Craig, 1988; Feder & Hofmann, 1999).

Functional enrichment analyses revealed tissue- and temperature-specific patterns of gene expression that appear to reflect functional priorities under thermal stress and align with the physiological demands of maintaining both survival and reproductive capacity.

Liver tissue showed the most differential expression (135 total DEGs across treatments) consistent with its many diverse roles in metabolism and detoxification during stress (Dou, et al., 2021). Heart tissue showed the weakest response (47 total DEGs across treatments), suggesting that maintenance of cardiac stability may take precedence over transcriptional flexibility (Drown et al., 2022). Brain tissue exhibited moderate differential expression (60 total DEGs across treatments), possibly balancing the need to maintain critical neural functions necessary to maintain courtship activity, continuous behavioral thermoregulation, and to perform risk assessment and make decisions to balance courtship effort with thermal risk. Lastly, testis tissue showed a moderately high transcriptional response (95 total DEGs across treatments), potentially reflecting the need to protect stored sperm from thermal damage while maintaining production of secretions essential for sperm transfer via formation of copulatory plugs (Friesen et al., 2013), as both of these functions are crucial for successful reproduction.

In brain tissue, heat stress resulted in a downregulation of genes associated with metabolic processes and RNA/DNA biosynthesis which may suggest a protective strategy to reduce energetically costly processes when body temperatures approach critical limits. The upregulation of structural functions and chromatin organization during cold stress may indicate activation of functional or structural plasticity. Interestingly, genes downregulated in response to cold stress were associated with metabolic processes, while genes simultaneously upregulated were associated with negative regulation of metabolic processes. Together, these opposing patterns of differential expression both appear to result in a suppression of metabolic processes. Thus, it appears that transcription and metabolism are not simply reduced when body temperatures drop, but transcription is actively increased to further suppress multiple metabolic processes.

Heart tissue showed the most conservative transcriptional response, with upregulation of protein catabolic processes under heat stress and translation processes under cold stress. This restrained response is consistent with the need to maintain cardiac stability and circulation during thermal stress, as cardiovascular function is essential for both thermoregulatory heat distribution and continued reproductive activity.

The liver exhibited the most extensive transcriptional plasticity with upregulation of metabolic, catabolic, and cellular signaling processes under heat stress, while simultaneously downregulating membrane organization and lipid transport. Under cold stress, the liver prioritized gene expression machinery and RNA processing. This broad transcriptional flexibility reflects the liver’s central role in metabolic homeostasis and detoxification during thermal stress, supporting the energetic demands of both thermoregulation and prolonged reproductive effort without feeding throughout the spring mating period.

Differential expression in testis tissue showed temperature-specific responses that appear to be directly relevant to reproductive function. Cold stress in testis tissue resulted in upregulation of genes involved in carbohydrate metabolism and translation, potentially supporting energy production and protein synthesis necessary for maintaining activity of sperm in cold temperatures. Heat stress resulted in downregulation of genes involved in cellular structure and sperm production. This suggests a possible protective mechanism to preserve gametes and reproductive tissues from thermal damage, or alternatively, it may simply be an indication of a non-adaptive, negative effect of heat stress on reproductive tissues.

These anticipatory tissue-specific transcriptional responses demonstrate that physiological plasticity operates as part of a coordinated system that enables male snakes to sustain courtship at body temperatures approaching behavioral and lethal thresholds and illustrates a finely tuned integration of molecular protection and behavioral risk-taking that simultaneously optimizes reproductive effort and survival in an extreme thermal environment.

### Phenotypic plasticity as the target of selection

Phenotypic plasticity itself is an evolved trait, and selection may act not only on mean trait values, but on the interaction of traits and environment across varying contexts to produce reaction norms that allow individuals to optimize fitness across variable environments (Via & Lande, 1985; Pigliucci, 2001; West-Eberhard, 2003; Dobzhansky, 1950).

The thermal biology of male *T. s. parietalis* is consistent with temperature-dependent reaction norms that allow individuals to dynamically balance the competing demands of survival and reproduction across different thermal and behavioral contexts. The consistent behavioral shifts observed at specific thermal thresholds (BT_vol_ and BT_court_) suggest that these are not simply physiological constraints but represent functional responses that could be subject to selection in unpredictable thermal environments.

The shift from courtship behavior to thermoregulation when body temperatures increase beyond BT_court_ is consistent with the framework of evolved priority rules that resolve conflicts between mutually exclusive phenotypic traits (McNamara et al., 1999, Lima & Dill, 1990). In this system, the priority rule governing male thermal behavior appears straightforward: males maintain courtship behavior until body temperature closely approaches the critical thermal maximum, at which point survival-oriented thermoregulation takes precedence.

This switching threshold is positioned such that males maximize reproductive effort while maintaining a narrow safety margin before reaching lethal temperatures. Males that switch to thermoregulatory behavior at lower body temperatures would sacrifice reproductive opportunities to competitors willing to accept greater thermal risk, whereas males that switch at higher temperatures would face increased mortality risk and sacrifice future reproductive potential entirely. The narrow margin between this switching threshold and lethal temperatures suggests either strong optimization, severe constraint, or both.

Furthermore, the universal upregulation of heat shock proteins across tissues and thermal treatments suggests that physiological plasticity may have evolved to synergistically support behavioral risk-taking rather than simply responding to thermal damage (Lindquist & Craig, 1988; Feder & Hoffman, 1999). This anticipatory molecular response appears to operate in concert with behavioral switching to enable males to sustain courtship near lethal temperatures. The timing of this response is critical, as an individual cannot wait for loss of motor control to occur before engaging in thermoregulatory behavior. By activating protective mechanisms at body temperatures well below the lethal maximum, transcriptional plasticity creates a molecular safety buffer that supports the risky behavioral decisions observed during courtship.

Whether this represents an anticipatory adaptation or a conserved stress response mechanism shared across ectotherms remains an open question. However, the consistency and timing we observed demonstrate functional coordination between physiological and behavioral plasticity.

### Plasticity buffering and possible effects on evolutionary adaptive potential

The extreme behavioral and physiological plasticity demonstrated by male garter snakes may create complex, paradoxical effects on evolutionary potential. Plasticity can buffer populations against environmental variation by allowing phenotypic adjustment without requiring genetic change potentially relaxing selection pressure on genetic variants that may otherwise act as the substrate of adaptive evolution (Price et al., 2003; Ghalambor et al., 2007). If plasticity enables individuals with diverse genotypes to succeed across a range of thermal conditions, this could maintain genetic variation while simultaneously reducing the immediate efficacy of selection on thermal tolerance traits. Thus, individuals capable of dynamic and appropriate adjustments to thermal behavior while concurrently mounting rapid transcriptional responses may succeed across a range of thermal conditions potentially reducing the benefits of genotypes that would otherwise confer an evolutionary advantage.

Although buffering effects may provide short-term population stability, it is also possible that it may constrain long-term adaptive potential if environmental conditions shift beyond the limits of plastic responses (Chevin et al., 2010). The very narrow margin between thermal-behavioral thresholds and the critical thermal maximum suggests that behavioral and physiological plasticity may already be approaching the boundaries of their evolutionary limits (Bennett et al., 2021). Thus, continued warming of the environment may soon exceed the capacity of plasticity to compensate, and thus directly expose genetic variation to selection (Gunderson & Stillman, 2015). Even so, phenotypic plasticity and the evolutionary capacity of genetic variation provide more adaptive flexibility than either mechanism alone (West-Eberhard, 2003). Whether current levels of genetic variation in thermal tolerance traits would be sufficient to respond to such selection remains unknown for this population.

### Conclusions

This study demonstrates that male red-sided garter snakes actively navigate an extreme thermal environment while maximizing reproductive opportunities. Our integrated approach reveals how behavioral and physiological plasticity operate simultaneously to balance survival and reproduction under severe thermal constraints to mitigate the effects of this major evolutionary trade-off.

Variable environments favor the evolution of plasticity over simple trait optima (West-Eberhard, 2003; Pigliucci, 2001). Thus, selective pressures act directly on reaction norms, shaping both the mean thermal responses and phenotypic state across varied thermal and reproductive contexts. These concepts are clearly demonstrated by male *T. s. parietalis* illustrating how multiple evolutionary processes interact to produce complex adaptive phenotypes. The precision of thermal decision-making, the speed of physiological responses, and the context-dependency of behavioral switching all suggest evolved solutions to the problem of balancing competing selective pressures. The result is a remarkably fine-tuned system that allows an individual to maximize reproductive effort while optimizing survival in a thermally dangerous environment.

## Data Availability

All sequence data is available in NCBI BioProject PRJNA1357357.

## Citations

1. Aleksiuk M and Gregory PT. (1974). Regulation of seasonal mating behavior in Thamnophis sirtalis parietalis. Copeia, 18, 681–689.

2. Aleksiuk, M., & Stewart, K. W. (1971). Seasonal Changes in the Body Composition of the Garter Snake (Thamnophis Sirtalis Parietalis) at Northern Latitudes. Ecology, 52(3). 10.2307/1937631

3. Amo, L., López, P., & Martín, J. (2004). Trade-offs in the choice of refuges by common wall lizards: Do thermal costs affect preferences for predator-free refuges? Canadian Journal of Zoology, 82(6). 10.1139/Z04-065

4. Andrews, S. (2010). FastQC - A quality control tool for high throughput sequence data. http://www.bioinformatics.babraham.ac.uk/projects/fastqc/. Babraham Bioinformatics.

5. Arak, A. (1983). Sexual selection by male-male competition in natterjack toad choruses. Nature, 306(5940). 10.1038/306261a0

6. Ardia, D. R., Cooper, C. B., & Dhondt, A. A. (2006). Warm temperatures lead to early onset of incubation, shorter incubation periods and greater hatching asynchrony in tree swallows Tachycineta bicolor at the extremes of their range. Journal of Avian Biology, 37(2). 10.1111/j.0908-8857.2006.03747.x

7. Ashburner, M., Ball, C. A., Blake, J. A., Botstein, D., Butler, H., Cherry, J. M., Davis, A. P., Dolinski, K., Dwight, S. S., Eppig, J. T., Harris, M. A., Hill, D. P., Issel-Tarver, L., Kasarskis, A., Lewis, S., Matese, J. C., Richardson, J. E., Ringwald, M., Rubin, G. M., & Sherlock, G. (2000). Gene ontology: Tool for the unification of biology. In Nature Genetics (Vol. 25, Issue 1). 10.1038/75556

8. Beever, E. A., Hall, L. E., Varner, J., Loosen, A. E., Dunham, J. B., Gahl, M. K., Smith, F. A., & Lawler, J. J. (2017). Behavioral flexibility as a mechanism for coping with climate change. In Frontiers in Ecology and the Environment (Vol. 15, Issue 6). 10.1002/fee.1502

9. Bellard, C., Bertelsmeier, C., Leadley, P., Thuiller, W., & Courchamp, F. (2012). Impacts of climate change on the future of biodiversity. In Ecology Letters (Vol. 15, Issue 4). 10.1111/j.1461-0248.2011.01736.x

10. Benjamini, Y., & Hochberg, Y. (1995). Controlling the False Discovery Rate: A Practical and Powerful Approach to Multiple Testing. Journal of the Royal Statistical Society: Series B (Methodological), 57(1). 10.1111/j.2517-6161.1995.tb02031.x

11. Bennett, J. M., Sunday, J., Calosi, P., Villalobos, F., Martínez, B., Molina-Venegas, R., Araújo, M. B., Algar, A. C., Clusella-Trullas, S., Hawkins, B. A., Keith, S. A., Kühn, I., Rahbek, C., Rodríguez, L., Singer, A., Morales-Castilla, I., & Olalla-Tárraga, M. Á. (2021). The evolution of critical thermal limits of life on Earth. Nature Communications, 12(1). 10.1038/s41467-021-21263-8

12. Besson, A. A., & Cree, A. (2010). A cold-adapted reptile becomes a more effective thermoregulator in a thermally challenging environment. Oecologia, 163(3). 10.1007/s00442-010-1571-y

13. Bogert, C. M. (1949). Thermoregulation in reptiles; a factor in evolution. Evolution; International Journal of Organic Evolution, 3(3). 10.1111/j.1558-5646.1949.tb00021.x

14. Bradbury, J. W., & Gibson, R. M. (1983). Leks and mate choice. In Mate Choice.

15. Bush, E., & Lemmen, D. (2019). Canada’s Changing Climate Report. In Environment and Climate Change Canada, Government of Canada.

16. Cease, A. J., Lutterschmidt, D. I., & Mason, R. T. (2007). Corticosterone and the transition from courtship behavior to dispersal in male red-sided garter snakes (Thamnophis sirtalis parietalis). General and Comparative Endocrinology, 150(1). 10.1016/j.ygcen.2006.07.022

17. Ceballos, G., Ehrlich, P. R., & Dirzo, R. (2017). Biological annihilation via the ongoing sixth mass extinction signaled by vertebrate population losses and declines. Proceedings of the National Academy of Sciences of the United States of America, 114(30). 10.1073/pnas.1704949114

18. Chevin, L. M., Lande, R., & Mace, G. M. (2010). Adaptation, plasticity, and extinction in a changing environment: Towards a predictive theory. PLoS Biology, 8(4). 10.1371/journal.pbio.1000357

19. Cowles, R. B. (1941). Observations on the Winter Activities of Desert Reptiles. Ecology, 22(2). 10.2307/1932207

20. Cowles, R. B., & Bogert, C. M. (1944). A preliminary study of the thermal requirements of desert reptiles. Bulletin of the American Museum of Natural History, 83.

21. Crews, D. (1984). Gamete production, sex hormone secretion, and mating behavior uncoupled. Hormones and Behavior, 18(1). 10.1016/0018-506X(84)90047-3

22. David, M., Dzamba, M., Lister, D., Ilie, L., & Brudno, M. (2011). SHRiMP2: Sensitive yet practical short read mapping. Bioinformatics, 27(7). 10.1093/bioinformatics/btr046

23. Dingemanse, N. J., & Wolf, M. (2010). Recent models for adaptive personality differences: A review. In Philosophical Transactions of the Royal Society B: Biological Sciences (Vol. 365, Issue 1560). 10.1098/rstb.2010.0221

24. Dobzhansky, T. (1950). Evolution in the tropics: Dobzhansky revisited. American Scientist, 38.

25. Dou, J., Cánovas, A., Brito, L. F., Yu, Y., Schenkel, F. S., & Wang, Y. (2021). Comprehensive RNA-Seq Profiling Reveals Temporal and Tissue-Specific Changes in Gene Expression in Sprague–Dawley Rats as Response to Heat Stress Challenges. Frontiers in Genetics, 12. 10.3389/fgene.2021.651979

26. Drown, M. K., Crawford, D. L., & Oleksiak, M. F. (2022). Transcriptomic analysis provides insights into molecular mechanisms of thermal physiology. BMC Genomics, 23(1). 10.1186/s12864-022-08653-y

27. Environment Canada. (2014). Canada’s sixth national report on climate change 2014 : actions to meet commitments under the United Nations Framework Convention on Climate Change.

28. Erisman, B., Heyman, W., Kobara, S., Ezer, T., Pittman, S., Aburto-Oropeza, O., & Nemeth, R. S. (2017). Fish spawning aggregations: where well-placed management actions can yield big benefits for fisheries and conservation. Fish and Fisheries, 18(1). 10.1111/faf.12132

29. Ernst, C. H., Creque, T. R., Orr, J. M., Hartsell, T. D., & Laemmerzahl, A. F. (2014). Operating body temperatures in a snake community of Northern Virginia. Northeastern Naturalist, 21(2). 10.1656/045.021.0205

30. Feder, M. E., & Hofmann, G. E. (1999). Heat-shock proteins, molecular chaperones, and the stress response: Evolutionary and ecological physiology. In Annual Review of Physiology (Vol. 61). 10.1146/annurev.physiol.61.1.243

31. Friesen, C. R., Powers, D. R., & Mason, R. T. (2017). Using whole-group metabolic rate and behaviour to assess the energetics of courtship in red-sided garter snakes. Animal Behaviour, 130. 10.1016/j.anbehav.2017.06.020

32. Friesen, C. R., Shine, R., Krohmer, R. W., & Mason, R. T. (2013). Not just a chastity belt: The functional significance of mating plugs in garter snakes, revisited. Biological Journal of the Linnean Society, 109(4). 10.1111/bij.12089

33. Gangloff, E. J., & Telemeco, R. S. (2018). High temperature, oxygen, and performance: Insights from reptiles and amphibians. Integrative and Comparative Biology, 58(1). 10.1093/icb/icy005

34. Garstka, W. R., Camazine, B., & Crews, D. (1982). Interactions of behavior and physiology during the annual reproductive cycle of the red-sided garter snake (Thamnophis sirtalis parietalis). Herpetologica, 38(1).

35. Gene Ontology Consortium. The Gene Ontology knowledgebase in 2023. Genetics. 2023 May 4;224(1). 10.1093/genetics/iyad031

36. Ghalambor, C. K., McKay, J. K., Carroll, S. P., & Reznick, D. N. (2007). Adaptive versus non-adaptive phenotypic plasticity and the potential for contemporary adaptation in new environments. Functional Ecology, 21(3). 10.1111/j.1365-2435.2007.01283.x

37. Gienapp, P., Teplitsky, C., Alho, J. S., Mills, J. A., & Merilä, J. (2008). Climate change and evolution: Disentangling environmental and genetic responses. In Molecular Ecology (Vol. 17, Issue 1). 10.1111/j.1365-294X.2007.03413.x

38. Glanville, E. J., & Seebacher, F. (2006). Compensation for environmental change by complementary shifts of thermal sensitivity and thermoregulatory behaviour in an ectotherm. Journal of Experimental Biology, 209(24). 10.1242/jeb.02585

39. Goto, S. G., & Kimura, M. T. (1998). Heat- and cold-shock responses and temperature adaptations in subtropical and temperate species of Drosophila. Journal of Insect Physiology, 44(12). 10.1016/S0022-1910(98)00101-2

40. Götz, S., García-Gómez, J. M., Terol, J., Williams, T. D., Nagaraj, S. H., Nueda, M. J., Robles, M., Talón, M., Dopazo, J., & Conesa, A. (2008). High-throughput functional annotation and data mining with the Blast2GO suite. Nucleic Acids Research, 36(10). 10.1093/nar/gkn176

41. Grabherr, M. G., Haas, B. J., Yassour, M., Levin, J. Z., Thompson, D. A., Amit, I., Adiconis, X., Fan, L., Raychowdhury, R., Zeng, Q., Chen, Z., Mauceli, E., Hacohen, N., Gnirke, A., Rhind, N., di Palma, F., Birren, B. W., Nusbaum, C., Lindblad-Toh, K., … Regev, A. (2011). Full-length transcriptome assembly from RNA-Seq data without a reference genome. Nature Biotechnology, 29(7). 10.1038/nbt.1883

42. Gregory, P. T. (1974). Patterns of spring emergence of the red-sided garter snake (Thamnophis sirtalis parietalis) in the Interlake region of Manitoba. Canadian Journal of Zoology, 52(8). 10.1139/z74-141

43. Gunderson, A. R., & Stillman, J. H. (2015). Plasticity in thermal tolerance has limited potential to buffer ectotherms from global warming. Proceedings of the Royal Society B: Biological Sciences, 282(1808). 10.1098/rspb.2015.0401

44. Haas, B. J., Papanicolaou, A., Yassour, M., Grabherr, M., Blood, P. D., Bowden, J., Couger, M. B., Eccles, D., Li, B., Lieber, M., Macmanes, M. D., Ott, M., Orvis, J., Pochet, N., Strozzi, F., Weeks, N., Westerman, R., William, T., Dewey, C. N., … Regev, A. (2013). De novo transcript sequence reconstruction from RNA-seq using the Trinity platform for reference generation and analysis. Nature Protocols, 8(8). 10.1038/nprot.2013.084

45. Howarth, C. J., & Ougham, H. J. (1993). Gene expression under temperature stress. New Phytologist, 125(1). 10.1111/j.1469-8137.1993.tb03862.x

46. Huey, R. B., Ma, L., Levy, O., & Kearney, M. R. (2021). Three questions about the eco-physiology of overwintering underground. Ecology Letters, 24(2). 10.1111/ele.13636

47. Huey, R. B., & Slatkin, M. (1976). Cost and benefits of lizard thermoregulation. The Quarterly Review of Biology, 51(3). 10.1086/409470

48. IPCC. (2021). IPCC: Summary for Policymakers. Climate Change 2021: The Physical Science Basis. Contribution of Working Group I to the Sixth Assessment Report of the Intergovernmental Panel on Climate Change.

49. Kearney, M., Shine, R., & Porter, W. P. (2009). The potential for behavioral thermoregulation to buffer “cold-blooded” animals against climate warming. Proceedings of the National Academy of Sciences of the United States of America, 106(10). 10.1073/pnas.0808913106

50. King, M. B., & Duvall, D. (1990). Prairie rattlesnake seasonal migrations: episodes of movement, vernal foraging and sex differences. Animal Behaviour, 39(5). 10.1016/S0003-3472(05)80957-1

51. Kingsolver, J. G., & Huey, R. B. (1998). Evolutionary analyses of morphological and physiological plasticity in thermally variable environments. American Zoologist, 38(3). 10.1093/icb/38.3.545

52. Kitchen, S. A., Crowder, C. M., Poole, A. Z., Weis, V. M., & Meyer, E. (2015). De novo assembly and characterization of four anthozoan (phylum Cnidaria) transcriptomes. G3: Genes, Genomes, Genetics, 5(11). 10.1534/g3.115.020164

53. Licht, P. (1964). The temperature dependence of myosin-adenosinetriphosphatase and alkaline phosphatase in lizards. Comparative Biochemistry And Physiology, 12(3). 10.1016/0010-406X(64)90063-5

54. Lima, S. L., & Dill, L. M. (1990). Behavioral decisions made under the risk of predation: a review and prospectus. Canadian Journal of Zoology, 68(4). 10.1139/z90-092

55. Lindquist, S., & Craig, E. A. (1988). THE HEAT-SHOCK PROTEINS. Annual Review of Genetics, 22(1), 631–677. 10.1146/annurev.ge.22.120188.003215

56. Liwanag, H. E. M. (2010). Energetic costs and thermoregulation in northern fur seal (Callorhinus ursinus) pups: The importance of behavioral strategies for thermal balance in furred marine mammals. Physiological and Biochemical Zoology, 83(6). 10.1086/656426

57. Lourdais, O., Bonnet, X., Shine, R., Denardo, D., Naulleau, G., & Guillon, M. (2002). Capital-breeding and reproductive effort in a variable environment: A longitudinal study of a viviparous snake. Journal of Animal Ecology, 71(3). 10.1046/j.1365-2656.2002.00612.x

58. Love, M. I., Huber, W., & Anders, S. (2014). Moderated estimation of fold change and dispersion for RNA-seq data with DESeq2. Genome Biology, 15(12). 10.1186/s13059-014-0550-8

59. Lutterschmidt, D. I., LeMaster, M. P., & Mason, R. T. (2006). Minimal overwintering temperatures of red-sided garter snakes (Thamnophis sirtalis parietalis): A possible cue for emergence? Canadian Journal of Zoology, 84(5). 10.1139/Z06-043

60. Lutterschmidt, W. I., & Hutchison, V. H. (1997). The critical thermal maximum: History and critique. In Canadian Journal of Zoology (Vol. 75, Issue 10). 10.1139/z97-783

61. Madsen, T., & Shine, R. (1996). Seasonal migration of predators and prey -A study of pythons and rats in tropical Australia. Ecology, 77(1). 10.2307/2265663

62. Masson-Delmotte, V. (2023). Climate change 2021 : the physical science basis : Working Group I contribution to the Sixth Assessment Report of the Intergovernmental Panel on Climate Change. 10.1017/9781009157889

63. McNamara, J. M., Gasson, C. E., & Houston, A. I. (1999). Incorporating rules for responding into evolutionary games. Nature, 401(6751). 10.1038/43869

64. Merilä, J., & Hendry, A. P. (2014). Climate change, adaptation, and phenotypic plasticity: The problem and the evidence. In Evolutionary Applications (Vol. 7, Issue 1). 10.1111/eva.12137

65. Meyer, E., Aglyamova, G. v., & Matz, M. v. (2011). Profiling gene expression responses of coral larvae (Acropora millepora) to elevated temperature and settlement inducers using a novel RNA-Seq procedure. Molecular Ecology, 20(17). 10.1111/j.1365-294X.2011.05205.x

66. Meyer, E., Aglyamova, G. v., Wang, S., Buchanan-Carter, J., Abrego, D., Colbourne, J. K., Willis, B. L., & Matz, M. v. (2009). Sequencing and de novo analysis of a coral larval transcriptome using 454 GSFlx. BMC Genomics, 10. 10.1186/1471-2164-10-219

67. Moore, I. T., Lemaster, M. P., & Mason, R. T. (2000). Behavioural and hormonal responses to capture stress in the male red-sided garter snake, Thamnophis sirtalis parietalis. Animal Behaviour, 59(3). 10.1006/anbe.1999.1344

68. Murphy-Klassen, H. M., Underwood, T. J., Sealy, S. G., & Czyrnyj, A. A. (2005). Long-term trends in spring arrival dates of migrant birds at delta marsh, Manitoba, in relation to climate change. Auk, 122(4). 10.1642/0004-8038(2005)122[1130:LTISAD]2.0.CO;2

69. O’Donnell, R. P., Shine, R., & Mason, R. T. (2004). Seasonal anorexia in the male red-sided garter snake, Thamnophis sirtalis parietalis. Behavioral Ecology and Sociobiology, 56(5). 10.1007/s00265-004-0801-x

70. Oostra, V., Saastamoinen, M., Zwaan, B. J., & Wheat, C. W. (2018). Strong phenotypic plasticity limits potential for evolutionary responses to climate change. Nature Communications, 9(1). 10.1038/s41467-018-03384-9

71. Parmesan, C. (2006). Ecological and evolutionary responses to recent climate change. In Annual Review of Ecology, Evolution, and Systematics (Vol. 37). 10.1146/annurev.ecolsys.37.091305.110100

72. Parmesan, C., & Yohe, G. (2003). A globally coherent fingerprint of climate change impacts across natural systems. Nature, 421(6918). 10.1038/nature01286

73. Pimm, S. L., Russell, G. J., Gittleman, J. L., & Brooks, T. M. (1995). The Future of Biodiversity. Science, 269(5222), 347–350. 10.1126/science.269.5222.347

74. Price, T. D., Qvarnström, A., & Irwin, D. E. (2003). The role of phenotypic plasticity in driving genetic evolution. In Proceedings of the Royal Society B: Biological Sciences (Vol. 270, Issue 1523). 10.1098/rspb.2003.2372

75. Ralpha Gibson, A., & Bruce Falls, B. (1979). Thermal biology of the common garter snake Thamnophis sirtalis (L.) -I. Temporal variation, environmental effects and sex differences. Oecologia, 43(1). 10.1007/BF00346674

76. Reznick, D., & Endler, J. A. (1982). The Impact of Predation on Life History Evolution in Trinidadian Guppies (Poecilia reticulata). Evolution, 36(1). 10.2307/2407978

77. Rumble, S. M., Lacroute, P., Dalca, A. v., Fiume, M., Sidow, A., & Brudno, M. (2009). SHRiMP: Accurate mapping of short color-space reads. PLoS Computational Biology, 5(5). 10.1371/journal.pcbi.1000386

78. Seebacher, F., & Franklin, C. E. (2005). Physiological mechanisms of thermoregulation in reptiles: A review. In Journal of Comparative Physiology B: Biochemical, Systemic, and Environmental Physiology (Vol. 175, Issue 8). 10.1007/s00360-005-0007-1

79. Seebacher, F., Tallis, J. A., & James, R. S. (2014). The cost of muscle power production: Muscle oxygen consumption per unit work increases at low temperatures in Xenopus laevis. Journal of Experimental Biology, 217(11). 10.1242/jeb.101147

80. Seebacher, F., White, C. R., & Franklin, C. E. (2015). Physiological plasticity increases resilience of ectothermic animals to climate change. Nature Climate Change, 5(1). 10.1038/nclimate2457

81. Shine, R., Harlow, P. S., Elphick, M. J., Olsson, M. M., & Mason, R. T. (2000). Conflicts between courtship and thermoregulation: The thermal ecology of amorous male garter snakes (Thamnophis sirtalis parietallis, Colubridae). Physiological and Biochemical Zoology, 73(4). 10.1086/317734

82. Shine, R., LeMaster, M. P., Moore, I. T., Olsson, M. M., & Mason, R. T. (2001). Bumpus in the snake den: Effects of sex, size, and body condition on mortality of red-sided garter snakes. Evolution, 55(3). 10.1111/j.0014-3820.2001.tb00792.x

83. Skinner, M., & Miller, N. (2020). Aggregation and social interaction in garter snakes (Thamnophis sirtalis sirtalis). Behavioral Ecology and Sociobiology, 74(5). 10.1007/s00265-020-2827-0

84. Somero, G. N. (1975). Temperature as a selective factor in protein evolution: The adaptational strategy of “compromise.” Journal of Experimental Zoology, 194(1). 10.1002/jez.1401940111

85. Somero, G. N. (1995). Proteins and temperature. In Annual Review of Physiology (Vol. 57). 10.1146/annurev.ph.57.030195.000355

86. Southwood, A., & Avens, L. (2010). Physiological, behavioral, and ecological aspects of migration in reptiles. In Journal of Comparative Physiology B: Biochemical, Systemic, and Environmental Physiology (Vol. 180, Issue 1). 10.1007/s00360-009-0415-8

87. Spellerberg, I. F. (1972). Thermal ecology of allopatric lizards (Sphenomorphus) in Southeast Australia -III. Behavioural aspects of thermoregulation. Oecologia, 11(1). 10.1007/BF00345706

88. Ultsch, G. R. (2006). The ecology of overwintering among turtles: Where turtles overwinter and its consequences. In Biological Reviews of the Cambridge Philosophical Society (Vol. 81, Issue 3). 10.1017/S1464793106007032

89. van de Ven, T. M. F. N., Fuller, A., & Clutton-Brock, T. H. (2020). Effects of climate change on pup growth and survival in a cooperative mammal, the meerkat. Functional Ecology, 34(1). 10.1111/1365-2435.13468

90. Via, S., & Lande, R. (1985). Genotype-Environment Interaction and the Evolution of Phenotypic Plasticity. Evolution, 39(3). 10.2307/2408649

91. Walther, G.-R., Post, E., Convey, P., Menzel, A., Parmesan, C., Beebee, T. J. C., Fromentin, J.-M., Hoegh-Guldberg, O., & Bairlein, F. (2002). Ecological responses to recent climate change. Nature, 416(6879), 389–395. 10.1038/416389a

92. West-Eberhard, M. J. (2003). Developmental Plasticity and Evolution. Oxford University Press.

93. Whittier, J. M., Mason, R. T., Crews, D., & Licht, P. (1987). Role of light and temperature in the regulation of reproduction in the red-sided garter snake, Thamnophis sirtalis parietalis. Canadian Journal of Zoology, 65(8). 10.1139/z87-320

94. Williams, C. M., Buckley, L. B., Sheldon, K. S., Vickers, M., Pörtner, H. O., Dowd, W. W., Gunderson, A. R., Marshall, K. E., & Stillman, J. H. (2016). Biological impacts of thermal extremes: mechanisms and costs of functional responses matter. Integrative and Comparative Biology, 56(1). 10.1093/icb/icw013

95. Williams, K. A., S. J. P., D. L. R., & D. C. R. (2025). Sitting in the open: how nest microclimate influences incubation behavior in an open-cup nesting passerine. Journal of Avian Biology, 2025(1), e03385.

96. Wright, R. M., Aglyamova, G. v., Meyer, E., & Matz, M. v. (2015). Gene expression associated with white syndromes in a reef building coral, Acropora hyacinthus. BMC Genomics, 16(1). 10.1186/s12864-015-1540-2

97. Zera, A. J., & Harshman, L. G. (2001). The physiology of life history trade-offs in animals. In Annual Review of Ecology and Systematics (Vol. 32). 10.1146/annurev.ecolsys.32.081501.114006

98. Zhao, F. Q. ing, Zhang, Z. W. ei, Qu, J. P. ing, Yao, H. D. ong, Li, M., Li, S., & Xu, S. W. en. (2014). Cold stress induces antioxidants and Hsps in chicken immune organs. Cell Stress & Chaperones, 19(5). 10.1007/s12192-013-0489-9

